# A transcription start site map in human pancreatic islets reveals functional regulatory signatures

**DOI:** 10.1101/812552

**Authors:** Arushi Varshney, Yasuhiro Kyono, Venkateswaran Ramamoorthi Elangovan, Collin Wang, Michael R. Erdos, Narisu Narisu, Ricardo D’Oliveira Albanus, Peter Orchard, Michael L. Stitzel, Francis S. Collins, Jacob O. Kitzman, Stephen C. J. Parker

## Abstract

Identifying the tissue-specific molecular signatures of active regulatory elements is critical to understand gene regulatory mechanisms. Here, we identify transcription start sites (TSS) using cap analysis of gene expression (CAGE) across 57 human pancreatic islet samples. We identify 9,954 reproducible CAGE tag clusters (TCs), ~20% of which are islet-specific and occur mostly distal to known gene TSSs. We integrated islet CAGE data with histone modification and chromatin accessibility profiles to identify epigenomic signatures of transcription initiation. Using a massively parallel reporter assay, we validate transcriptional enhancer activity (5% FDR) for 2,279 of 3,378 (~68%) tested islet CAGE elements. TCs within accessible enhancers show higher enrichment to overlap type 2 diabetes genome-wide association study (GWAS) signals than existing islet annotations, which emphasizes the utility of mapping CAGE profiles in disease-relevant tissue. This work provides a high-resolution map of transcriptional initiation in human pancreatic islets with utility for dissecting functional enhancers at GWAS loci.

## Introduction

Genome wide association studies (GWAS) for complex diseases such as type 2 diabetes (T2D) have identified hundreds of signals associated with disease risk, however, most of these lie in non protein-coding regions and the underlying mechanisms are still unclear (Mahajan *et al*. 2018). T2D GWAS variants are highly enriched to overlap islet-specific enhancer regions, which suggests these variants affect gene expression (Parker *et al*. 2013; Pasquali *et al*. 2014; Quang *et al*. 2015; Thurner *et al*. 2018; Cebola 2019). Many GWAS signals are marked by numerous SNPs in high linkage disequilibrium (LD) which makes identifying causal SNP(s) extremely difficult using genetic information alone.

To delineate regulatory elements, profiling histone modifications such as the enhancer associated H3 lysine 27 acetylation (H3K27ac) (Creyghton *et al*. 2010; Zhou *et al*. 2011), and the promoter associated H3 lysine 4 trimethylation (H3K4me3) (Mikkelsen *et al*. 2007; Adli *et al*. 2010; Zhou *et al*. 2011) among others can be useful. However, these methods identify genomic regions typically spanning hundreds of base pairs. Profiling TF-accessible chromatin regions can identify the functional DNA bases with these broad regulatory elements. Such approaches in pancreatic islets have nominated causal gene regulatory mechanisms (Fadista *et al*. 2014; Bunt *et al*. 2015; Varshney *et al*. 2017; Roman *et al*. 2017; Thurner *et al*. 2018; Mahajan *et al*. 2018; Rai *et al*.2019). Studies have shown that transcription is a robust predictor of enhancer activity and a subset of enhancers are transcribed into enhancer RNA (eRNA) (Andersson *et al*. 2014; Mikhaylichenko *et al*. 2018). eRNAs are nuclear, short, mostly-unspliced, 5’ capped, usually non-polyadenylated and usually bidirectionally transcribed (Kim *et al*. 2010; Melgar *et al*. 2011; Andersson *et al*. 2014). Therefore, identifying the location of transcription initiation can pinpoint active enhancer regulatory elements in addition to active promoters.

Genome-wide sequencing of 5’ capped RNAs using CAGE can detect TSSs (Kim *et al*. 2010; Andersson *et al*. 2014). CAGE can be applied on RNA samples from hard to acquire biological tissue such as islets and does not require live cells that are imperative for other TSS profiling techniques such as GRO-cap seq (Core *et al*. 2008, 2014; Lopes *et al*. 2017). The functional annotation of the mammalian genome (FANTOM) project (The FANTOM Consortium *et al*. 2014) has generated an exhaustive CAGE expression atlas across 573 primary cell types and tissues, including the pancreas. However, pancreatic islets that secrete insulin and are relevant for T2D and related traits, constitute only ~1% of the pancreas tissue. Therefore, the pancreas TSS map does not accurately represent the islet TSS landscape. To date, there are no publicly-available CAGE datasets for islet tissue. Motivated by these reasons, here we profiled the islet transcriptome using CAGE and present a TSS map of pancreatic islets, validate this map using a massively parallel reporter assay (MPRA), and perform integrative analyses across existing epigenomic datasets to reveal molecular signatures of non-coding islet elements and their role in T2D and related traits.

## Results

### The CAGE landscape in human pancreatic islets

We analyzed transcriptomes in 71 human pancreatic islet total RNA samples obtained from unrelated organ donors by performing CAGE according to the no-amplification non-tagging CAGE libraries for Illumina sequencers (nAnT-iCAGE) protocol followed by sequencing (Murata *et al*. 2014) (see Methods). After data processing and quality control (QC) measures, we selected 57 pass-QC samples and identified CAGE tag clusters (TCs) in each islet sample in a strand-specific manner using the paraclu method (Frith *et al*. 2008). We then identified a ‘consensus’ set of reproducible islet TCs by merging TCs on each strand across samples and retained segments supported by at least 10 samples (Supplementary Figure 1). This conservative consensus set included 9,954 TCs with median length of 176 bp (Supplementary Figure 2), spanning a total genomic territory of ~2.4 Mb. As a resource to identify transcribed gene isoforms in islets, Supplementary table 1 includes the closest identified islet TCs to known gene TSSs (Gencode V19, (Harrow *et al*. 2012)). To explore the chromatin landscape underlying islet TCs, we overlayed publicly-available ChIP-seq data for five histone modifications integrated into 11 distinct chromatin states using ChromHMM (Ernst and Kellis 2010, 2012; Ernst *et al*. 2011) (Supplementary Figure 3, see Methods), along with bulk and single nucleus ATAC-seq data in islets (Varshney *et al*. 2017; Rai *et al*. 2019). Figure 1A shows an example islet TC in the intronic region of the *ST18* gene that overlaps an islet active TSS chromatin state and an ATAC-seq peak. Importantly, this region does not overlap any annotated TSS using conservative definitions from coding/non-coding/pseudogene genes in both Gencode V19, the official hg19 release, and Gencode V33 lifted over to hg19 (V33lift37). The regulatory activity of this element was validated by the VISTA project in an *in vivo* reporter assay in mouse embryos (Visel *et al*. 2007), which further confirms the transcriptional enhancer capacity of this islet intronic TC..

**Figure 1:**
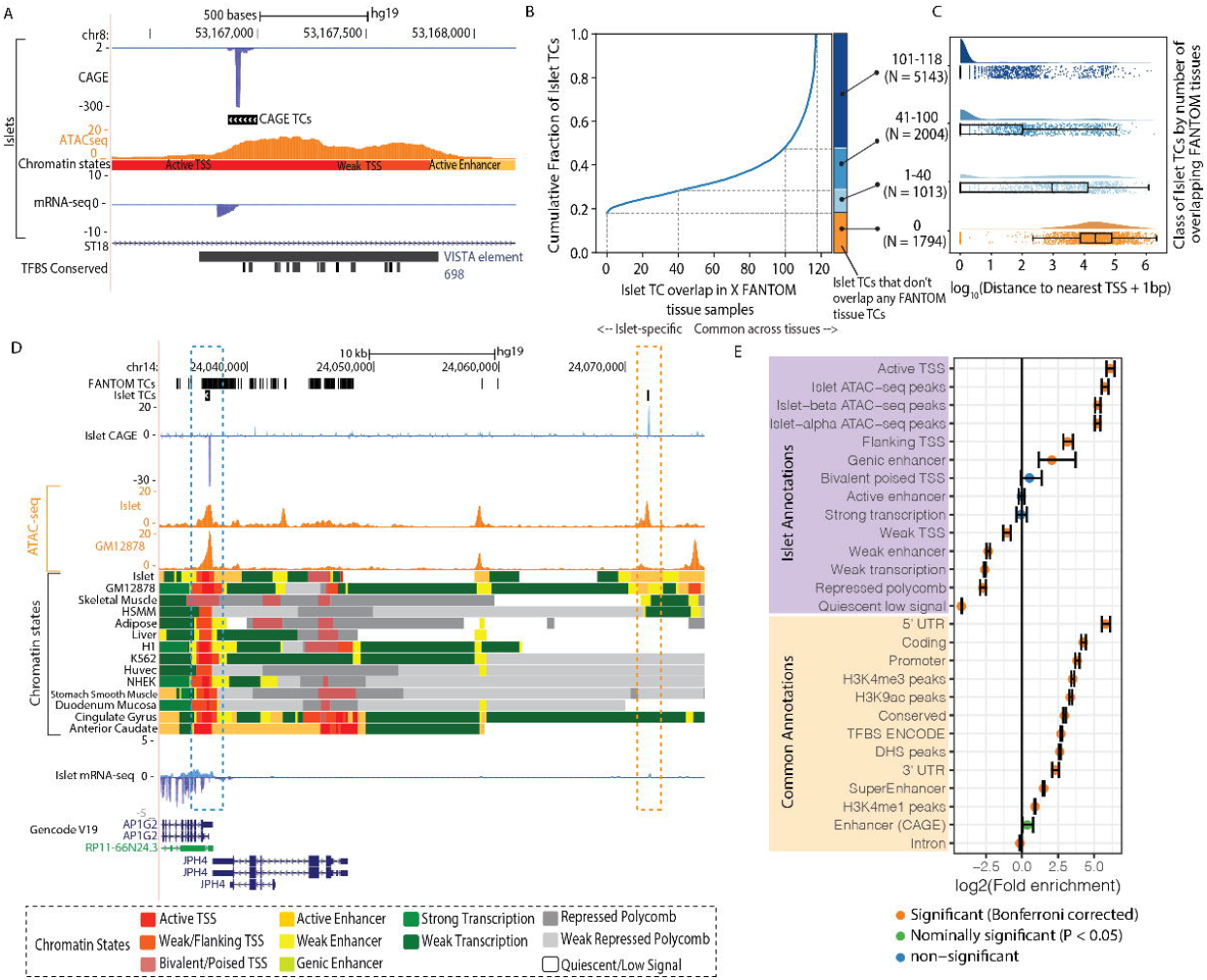
Islet CAGE tag cluster identification. A: Genome browser view of the intronic region of the *ST18* gene as an example locus where an islet TC overlaps an islet ATAC-seq peak and active TSS chromatin state. This TC also overlaps an enhancer element which was validated by the VISTA project (Visel *et al*. 2007). Also shown is the human-mouse-rat conserved TF binding site (TFBS) track from the Transfac Matrix Database (Matys *et al*. 2006). B: Cumulative fraction of islet TC segments overlapping with TCs identified in x FANTOM tissues. C: Distribution of the log10(distance to the nearest known protein coding gene TSS + 1bp) while classifying Islet TC segments by the number of FANTOM tissues where TCs overlap. Number of TC segments in each category are shown in parentheses. D: Genome browser view of an example locus near the *AP1G2* gene that highlights an islet TC (blue box) that is also identified in FANTOM tissues (FANTOM TCs track is a dense depiction of TCs called across 118 human tissues), occurs in a ATAC-seq peak region in both islets and lymphoblastoid cell line GM12878 (ATAC-seq track) and overlaps active TSS chromatin states across numerous tissues. Another islet TC (orange box) ~34 kb distal to the *AP1G2* gene is not identified as a TC in other FANTOM tissues, occurs in an islet ATAC-seq peak and a more islet-specific active enhancer chromatin state region. E: Enrichment of islet TCs to overlap islet chromatin state and other common annotations. Error bars represent the 95% confidence intervals. Colors represent significant enrichment after Bonferroni correction accounting for 40 total annotations (see Methods for references of common annotations, Supplementary Table 2), nominal enrichment (P value < 0.05) or non-significant enrichment.

We next compared islet CAGE data with CAGE data available in 118 human tissues through the FANTOM project (The FANTOM Consortium *et al*. 2014). We identified TCs in 118 FANTOM human tissues using the same approach we used on our islets. To gauge tissue-specificity, we calculated for each islet TC segment the number of FANTOM tissues in which TCs overlapped. We observed that ~20% of islet TCs were unique to islets (1,974 TCs had no overlap with TCs in any FANTOM tissues), whereas about ~60% of islet TCs were shared across 60 or more FANTOM tissues (Figure 1B). We then asked if the more islet-specific TCs occurred more distal to known protein coding gene TSSs. Categorizing islet TC segments by the number of FANTOM tissues in which they overlap TCs (colored bars in Figure 1B), we observed that islet-specific TCs (no overlap with FANTOM) occurred farthest from annotated TSSs, and TCs overlapping more FANTOM tissues occurred closer to annotated TSSs (Figure 1C). We highlight an example locus where an islet TC in the *AP1G2* gene occurs in active TSS chromatin states across multiple tissues, and overlaps shared ATAC-seq peaks in islet and the lymphoblastoid cell line GM 12878 (Buenrostro *et al*. 2013) (Figure 1D, blue box). TCs across FANTOM tissues are identified in this region (Figure 1D, FANTOM TCs track). The islet TC segment (Figure 1D, blue dashed box) overlaps TCs in 88 FANTOM tissues. Another islet TC ~34kb away, however, occurs in a region lacking gene annotations, and overlaps a more islet-specific active enhancer chromatin state and ATAC-seq peak (Figure 1D, orange dashed box). This region was not identified as a TC in any of the 118 analyzed FANTOM tissues. At other islet-relevant loci such as the Potassium Channel Subfamily K gene *KCNK16* TSS, we observe TCs in islets but not in other FANTOM tissues including the whole Pancreas (Supplementary Figure 4). Collectively, these results highlight that CAGE profiling in islets identifies islet-specific sites of transcription initiation.

We next asked if islet TCs preferentially overlap certain genomic annotations. We computed the enrichment of islet TCs to overlap islet annotations such as active TSS, enhancer and other chromatin states (see Methods) and ATAC-seq peaks in bulk islets and in islet alpha and beta cells (Varshney *et al*. 2017; Rai *et al*. 2019). We also included ‘common’ annotations such as known gene promoters, coding, untranslated regions (UTR), or annotations such as super enhancers, or histone-modification ChIP-seq peaks that were aggregated across multiple cell types. We observed that islet TCs were highly enriched to overlap islet active TSS chromatin states (fold enrichment = 69.72, P value = 0.0001, Figure 1E, Supplementary Table 2). This result is expected since CAGE profiles transcription start sites where the underlying chromatin is more likely to resemble the ‘active TSS’ chromatin state. TCs were also enriched to overlap bulk islet and islet alpha and beta cell ATAC-seq peaks (for all three annotations, fold enrichment > 37.58, P value = 0.0001, Figure 1E), signifying that the identified transcription initiation sites constitute TF-accessible chromatin.

### Integrating CAGE TCs with epigenomic information

We further explored CAGE profiles relative to the underlying chromatin landscape to identify signatures of transcription initiation. We first overlayed CAGE profiles over ATAC-seq data. Aggregated CAGE signal over ATAC-seq narrow peak summits highlighted a bidirectional pattern of transcription initiation flanking the ATAC-seq peak summit on both strands (Figure 2A). Conversely, on anchoring the ATAC-seq signal over islet TC centers we observed that the summit of the ATAC-seq signal lies upstream of the TC center (Figure 2B). In these analyses, upstream or downstream positions were determined from the strand the CAGE signal mapped to. We next asked if TF binding sites were more enriched to occur upstream or downstream of the TC. We utilized TF footprint motifs previously identified using islet ATAC-seq data and TF DNA binding position weight matrices (PWMs) (Wang *et al*. 2018)These footprint motifs represent putative TF binding sites that are also supported by accessible chromatin profiles, as opposed to TF motif matches that are only informed by DNA sequence. We observe that most TF footprint motifs were more enriched to overlap the 500 bp TC upstream region compared to the 500 bp downstream region relative to TCs (Figure 2C). These observations show that, as expected, the region just upstream of the TC is highly accessible where more TF binding events occur, and indicate the high quality of our islet TC map.

**Figure 2.**
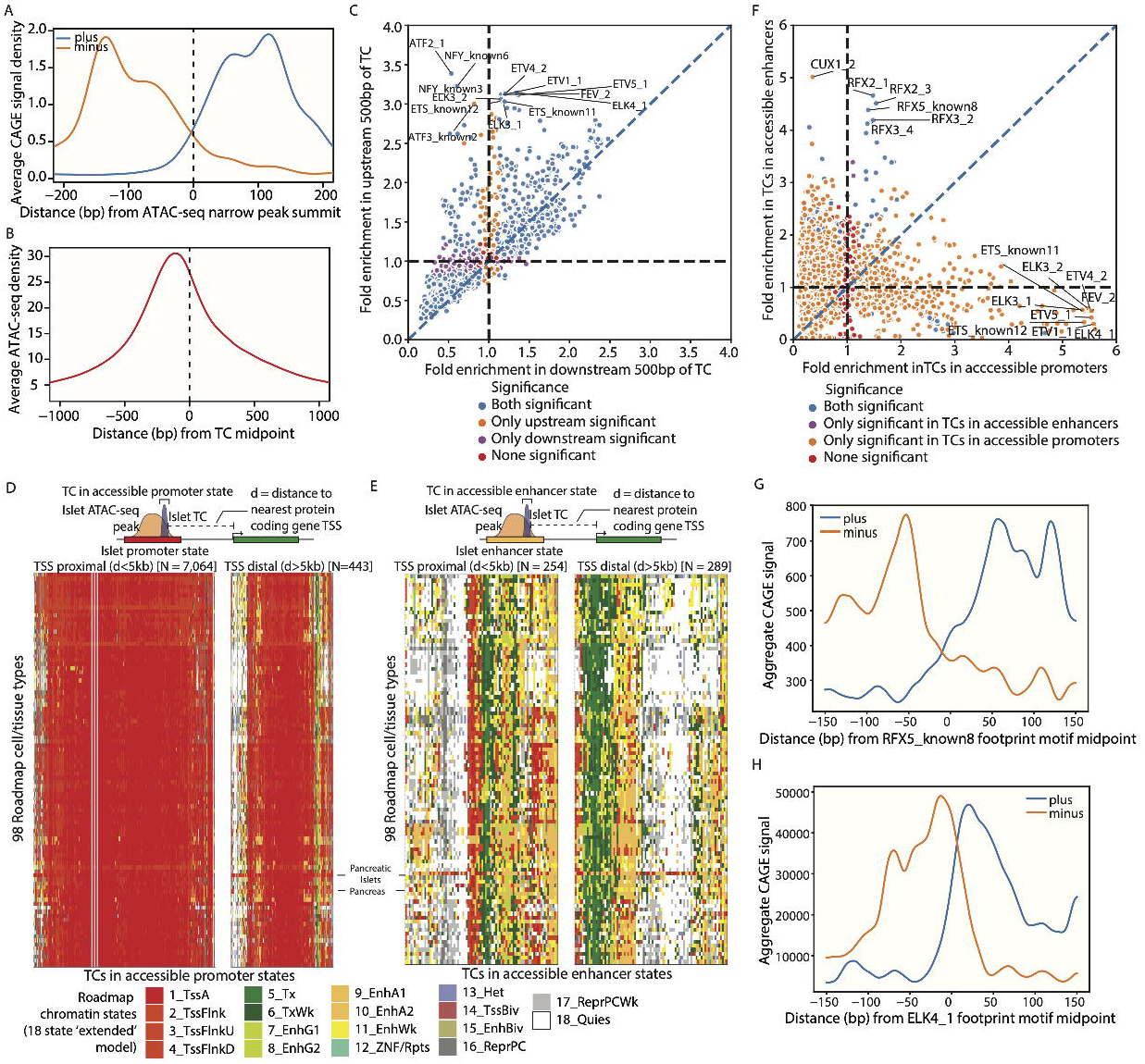
Integrating Islet CAGE TCs with other epigenomic information reveals characteristics of transcription initiation. A: Aggregate CAGE profiles over ATAC-seq peak summits. B: Aggregate ATAC-seq profile over TC midpoints. C: Enrichment of TF footprint motifs to overlap 500 bp upstream region (y axis) vs 500 bp downstream region (x axis) of islet TCs. Colors denote if a TF footprint motif was significantly enriched (5% FDR correction, Benjajmini-Yekutieli method) to overlap only upstream regions, only downstream regions, both or none. D: Chromatin state annotations across 98 Roadmap Epigenomics cell types (using the 18 state ‘extended model’ (The Roadmap Epigenomics Consortium *et al*. 2015) for TC segments that occur in islet promoter chromatin states and overlap ATAC-seq peaks. These segments were segregated into those occurring 5kb proximal (left, N=7,064 TC segments) and distal (right, N=443 TC segments) to known protein coding gene TSS (Gencode V19). E: Chromatin state annotations across 98 Roadmap Epigenomics cell types for TC segments that occur in islet enhancer chromatin states and overlap ATAC-seq peaks, segregated into those occurring 5kb proximal (left, N=254 TC segments) and distal (right, N=289 TC segments) to known protein coding gene TSS. F: Enrichment of TF footprint motifs to overlap TCs occurring in accessible enhancer chromatin states (y axis) vs TCs occurring in accessible promoter chromatin states (x axis). Colors denote if a TF footprint motif was significantly enriched (5% FDR correction, Benjajmini-Yekutieli method) to overlap only TCs in accessible enhancer regions, only TCs in accessible promoter regions, both or none. G: Aggregate CAGE profiles centered and oriented relative to RFX5_known8 footprint motifs occurring in 5kb TSS distal regions. H: Aggregate CAGE profiles centered and oriented relative to ELK4_1 footprint motifs.

We next explored the characteristics of TCs that occur in the two main regulatory classes - promoter and enhancers, relative to each other. We focussed on transcribed, accessible regions in promoter or enhancer chromatin states - namely TCs overlapping ATAC-seq peaks in promoter (active, weak or flanking TSS) chromatin states or enhancer (active, weak or genic enhancer) chromatin states. We considered the proximity of these elements to known gene TSSs and further classified the segments as TSS proximal or distal using a 5kb distance threshold from the nearest protein coding genes (Gencode V19). We then explored the chromatin landscape at these regions across 98 Roadmap Epigenomics cell types for which chromatin state annotations are publicly available (18 state ‘extended model’, see Methods) (The Roadmap Epigenomics Consortium *et al*.2015). We observed that TSS proximal islet TCs in accessible islet promoter chromatin states (N = 7,064 segments) were nearly ubiquitously identified as promoter chromatin states across roadmap cell types (Figure 2D, left). A subset of TSS distal islet TCs in accessible islet promoter chromatin states (N = 443 segments) were more specific for pancreatic islets (Figure 2D, right). In contrast, we observed that islet TCs in accessible islet enhancer chromatin states, both proximal (N = 254 segments) and distal (N = 289 segments) to known gene TSSs were more specifically identified as enhancer states in pancreatic islets (Figure 2E). This pattern was more clear for Roadmap pancreatic islet segmentations compared to the whole pancreas segmentations (Figure 2D and E, labelled) which further emphasizes the important differences between that our islet TC profiles highlight in pancreatic islets compared to whole pancreas tissue.

Having observed differences in cell type-specificities in islet TCs in promoter vs enhancer states, we next asked if TFs displayed preferences to bind in these regions. We observed that footprint motifs for the regulatory factor X (RFX) TF family were highly enriched (for 5 different motifs, >3 fold, P value = 0.0001) in TCs in accessible enhancers (Figure 2F). On the other hand, TCs in accessible promoter regions were highly enriched to overlap footprint motifs of the E26 transformation-specific (ETS) TF family (Figure 2F). We observe divergent aggregate CAGE profiles over TF footprint motifs enriched in enhancers; for example RFX5_known8 footprint motifs in 5kb TSS distal regions and ELK4_1 motif (Figure 2G and H). Taken together, these results show the differences in the characteristics of transcription initiation sites based on the underlying chromatin context.

### Experimental validation of transcribed regions

We next sought to experimentally validate the transcriptional activity of islet CAGE-profiled regions. We utilized an MPRA platform wherein thousands of elements can be simultaneously tested by including unique barcode sequences for each element and determining the transcriptional regulatory activity using sequencing-based barcode quantification (Melnikov *et al*. 2012; Arnold *et al*. 2013; Neumayr *et al*. 2019). We generated a library of 7,188 CAGE elements (198 bp each, see Methods) and cloned these sequences into the MPRA vector 3’ to the polyA signal of the GFP gene, which contained a 16 bp barcode. Our experimental setup therefore assays the ‘enhancer’ activity of the CAGE element where the readout is the transcript levels of the GFP-embedded barcode sequences corresponding to the CAGE element. We transfected the MPRA libraries into the rat beta cell insulinoma (INS1 832/13) cell line in triplicate, extracted DNA and RNA and sequenced the barcodes. We added 6 bp unique molecular identifier (UMI) sequences before the PCR amplification of the RNA libraries to enable accounting for PCR duplicates while quantifying true biological RNA copies. We selected barcodes that were observed with at least 10 DNA counts and non-zero RNA counts in at least one replicate, and selected CAGE elements that were observed with at least two such qualifying barcodes. This filtering procedure resulted in 3,378 CAGE elements. We observed high correlations between the normalized sum of RNA counts of the CAGE element barcodes across the three biological replicates (Pearson r = 0.97, Supplementary Figure 5). We quantified the enhancer reporter activity of the 3,378 CAGE elements by modeling the RNA and DNA barcode counts using generalized linear models (GLMs) implemented in the MPRAnalyze package (Ashuach *et al*. 2019) (Supplementary table 3). We then tested for significant transcriptional activity against a null model (see Methods). We observed that ~68% (N = 2,279) of the testable CAGE elements showed significant regulatory activity (5% FDR) (Figure 3A, top). After classifying CAGE elements based on the underlying chromatin landscape such as - promoter (active, weak or flanking TSS) chromatin state, enhancer (active, weak or genic enhancer) chromatin state or other chromatin state overlap in islets, we observed that a larger fraction of CAGE elements overlapping the promoter chromatin states had significant transcriptional activity compared to elements overlapping enhancer states, which was in turn higher than CAGE elements in other chromatin states (Figure 3A, bottom). We also observed that the CAGE elements in promoter chromatin states had higher MPRA activity Z scores compared to the elements in enhancer chromatin states (Wilcoxon rank sum test P = 1.02×10^-6^) (Figure 3B). Z scores for the CAGE elements that overlapped ATAC-seq peaks were significantly higher than the elements that did not occur in peaks (Wilcoxon rank sum test P = 5.50×10^-16^) (Figure 3C). Z scores for CAGE elements 5kb proximal to protein-coding gene TSSs (Gencode V19) were higher than CAGE elements that were distal to gene TSS locations (Wilcoxon rank sum test P = 5.38×10^-9^) (Supplementary Figure 6). These results highlight that TC elements in accessible promoter chromatin states show higher enhancer reporter activities, and is consistent with a recent MPRA study that used the GM12878 lymphoblastoid cell line (Wang *et al*. 2018).

**Figure 3.**
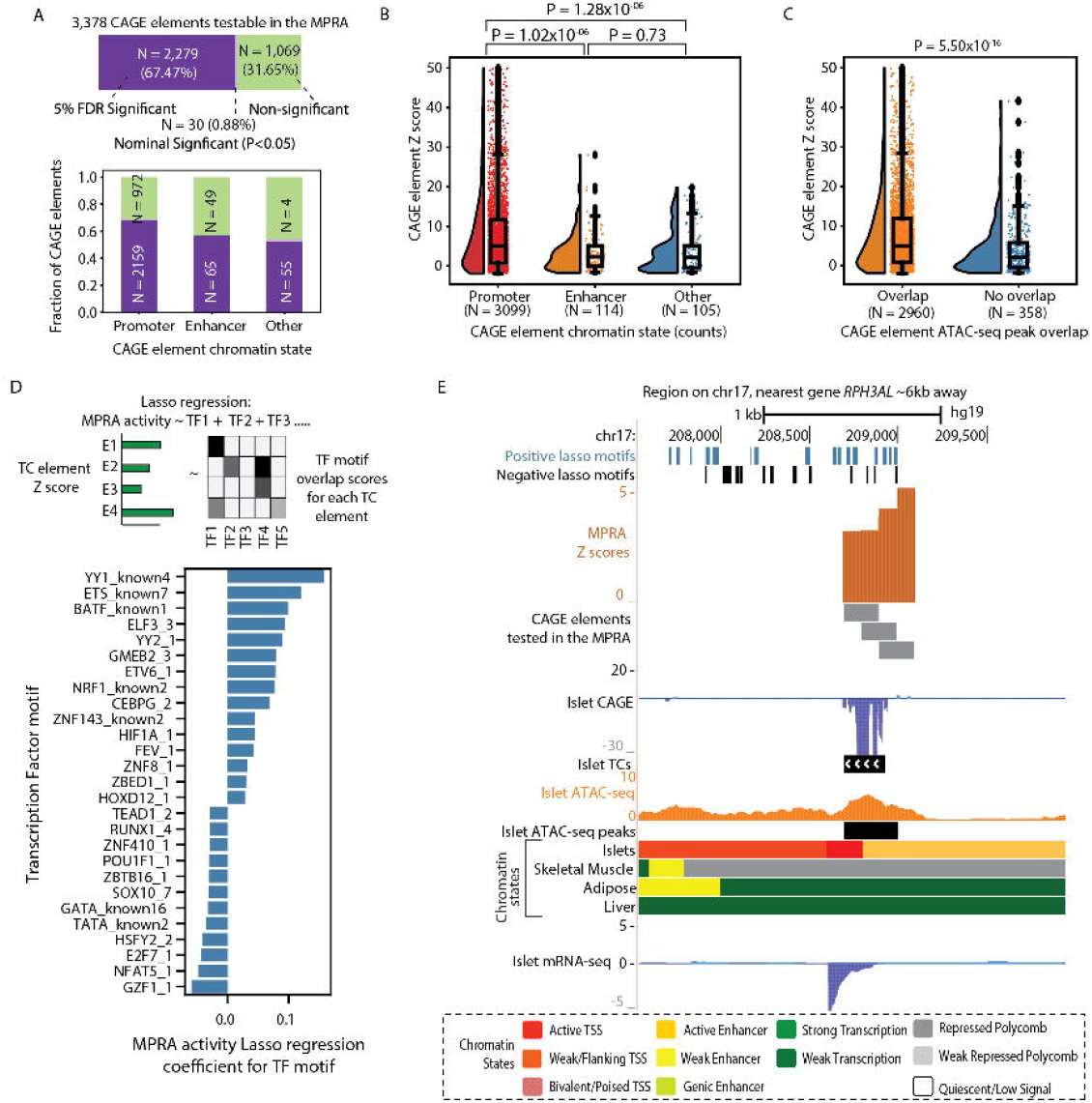
Experimental validation of CAGE elements using MPRA assay: A: (Top) Number and fraction of CAGE elements that show significant (5% FDR), nominal (P < 0.05) or non-significant transcriptional activity in the MPRA assay performed in rat beta cell insulinoma (INS1 832/13) cell line model. (Bottom) Proportion of CAGE elements overlapping promoter (active, weak or flanking TSS), enhancer (active, weak or genic enhancer) or other chromatin states that showed significant transcriptional activity in the MPRA assay. B. MPRA activity Z scores for CAGE elements overlapping in promoter, enhancer or other chromatin states. C: MPRA activity Z scores for CAGE elements that overlap ATAC-seq peak vs CAGE elements that do not overlap peaks. D: (Top) An overview of the lasso regression model to predict the MPRA activity Z scores of CAGE elements as a function of the TF motif scan scores within the element. (Bottom) Top 30 TF motifs with non-zero coefficients from the model. E: An example locus on chr17, where the nearest gene *RPH3AL* lies ~6kb away, an islet TC overlaps active TSS and enhancer chromatin states and an ATAC-seq peak. Elements overlapping this TC showed significant transcriptional activity in the MPRA assay. The CAGE profile coincides with islet mRNA profile that is detected despite no known gene annotation in the region and the nearest protein coding gene is ~6kb away. Also shown are occurrences of TF motifs with positive or negative lasso regression coefficients from the analysis in D.

We then asked if sequence-based features of the CAGE elements such as the occurrence of TF motifs could predict the activities of these elements in the MPRA assay. We used linear regression to model transcriptional activity as a function of TF motif overlap. Since thousands of TF motifs are available and many are correlated with each other, it is useful to identify a subset of more relevant motifs. For this purpose, we utilized the least absolute shrinkage and selection operator (LASSO) procedure which uses shrinkage, constraining the model parameters such that some regression coefficients are shrunk to zero. This results in a simpler model with a subset of selected features. We trained the LASSO regression model with the CAGE element MPRA Z scores as the response variable and the TF motif scan scores for 540 representative TF motifs in each CAGE element as predictors (see Methods). We report the TF motifs with the top 30 lasso regression coefficients (Figure 3D, full results in Supplementary Table 4). We observed that TF motifs from the ETS family showed positive lasso coefficients, indicating that these sequence elements are associated with high transcriptional activity. Indeed, we earlier observed that these motifs were also highly enriched to occur in TCs in accessible promoter chromatin state regions (Figure 2E). Other TF motifs with positive lasso coefficients included CEBP, YY1, and NRF-1 (Figure 3D). NRF-1, for instance, has been identified to be relevant for islet biology, as its target genes are downregulated in diabetic individuals (Patti *et al*. 2003); knockdown of this gene in the mouse insulinoma cell line (MIN6) and beta cell specific Nrf1-knockout mice resulted in impaired glucose responsiveness, elevated basal insulin release and decreased glucose-stimulated insulin secretion (GSIS) (Zheng *et al*. 2015).

In Figure 3E, we highlight an islet TC for which we tested three tiled elements (Figure 3E, MPRA elements track), which occurred in active TSS and enhancer states in islets and overlapped an islet ATAC-seq peak. All three elements showed significant transcriptional activity in our assay (Z score > 2.94, P values < 0.001). The active elements overlapped more TF motifs with positive lasso regression coefficients (Figure 3E, positive lasso motifs track) compared to TF motifs with negative lasso regression coefficients (Figure 3E, negative lasso motifs track), consistent with the observed active enhancer signal from the MPRA.

Through these analyses we experimentally validated ~68% of testable CAGE elements for significant transcriptional regulatory activity in a rodent islet beta cell model system, and identified TF motifs associated with high transcriptional activities.

### CAGE profiles augment functional genomic annotations to better fine-map GWAS loci

Observing the molecular signature of islet TCs in different epigenomic contexts and validating the activities of these elements, we next asked if islet TCs taken as an additional layer of functional genomic information could supplement our understanding of GWAS or islet expression quantitative loci (eQTL) associations. We classified genomic annotations based on layers of epigenomic data such as a) histone modification based chromatin states, b) accessible regions within the chromatin states and c) transcribed and accessible regions within the chromatin states. We then computed enrichment for T2D GWAS loci (Mahajan *et al*. 2018) to overlap these annotations using a Bayesian hierarchical model implemented in the fGWAS tool (Pickrell 2014). This method utilizes not only the genome wide significant loci but leverages full genome wide association summary statistics such that marginal associations can also be accounted for. We observed that TCs in accessible enhancer regions were the most highly enriched for T2D GWAS signals among the annotations tested (Figure 4A, left). We also computed enrichment for annotations to overlap islet eQTL (Varshney *et al*. 2017) and observed that TCs in accessible regions in both enhancers and promoters were the most highly enriched (Figure 4A, right). These data suggest that including TC information with other functional genomics data help illuminate relevant regions for the genetic control of gene expression and trait association signals.

**Figure 4.**
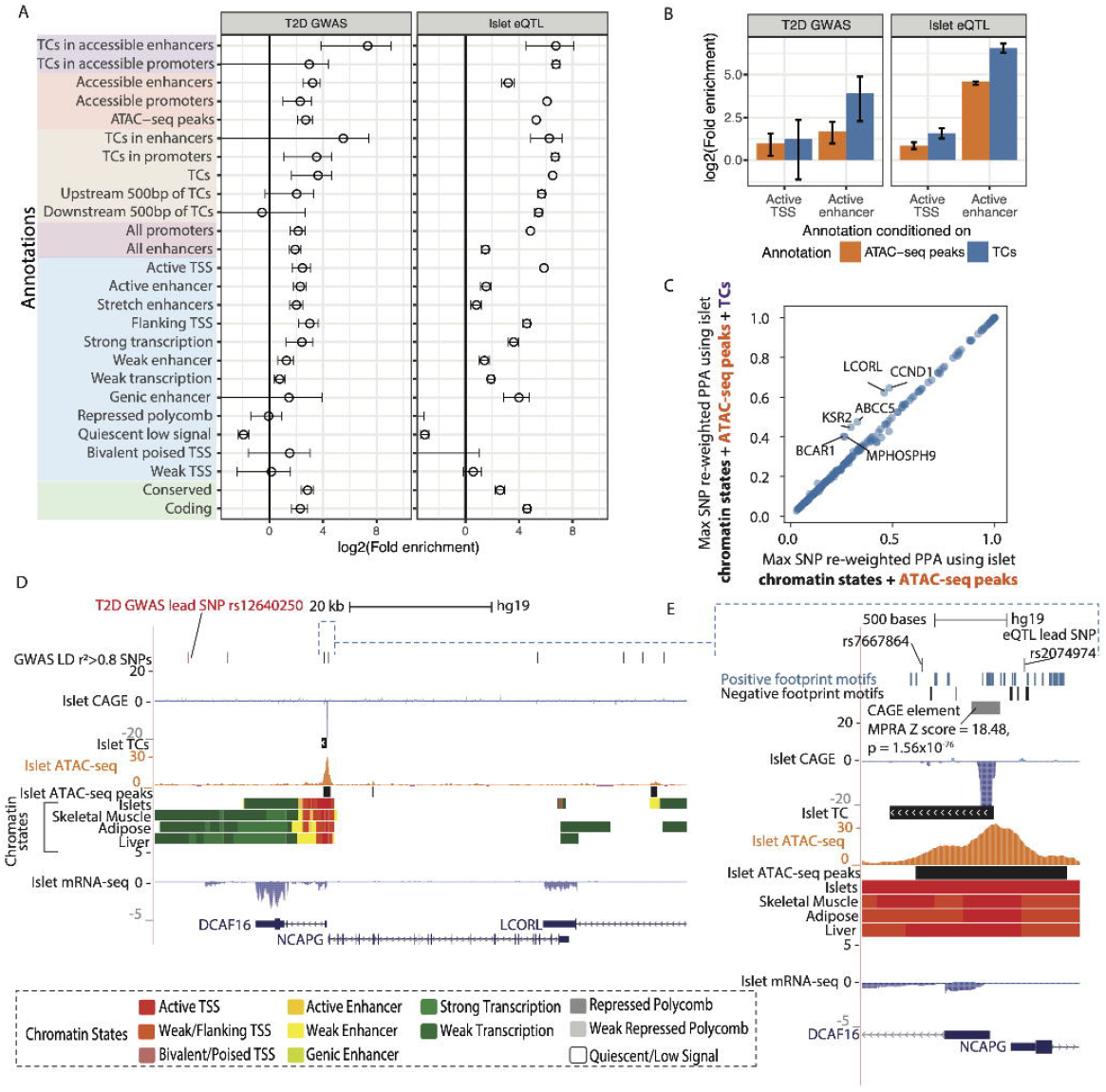
Islet TCs supplement functional understanding of GWAS and eQTL associations and help nominate causal variants. A: Enrichment of T2D GWAS (left) or islet eQTL (right) loci in annotations that comprise different levels of epigenomic information, including chromatin state, ATAC-seq and TCs. Annotations defined using combinations of these datasets are depicted with different colors on the y axis. Enrichment was calculated using fGWAS (Pickrell 2014) using summary statistics from GWAS (left) (Mahajan *et al*. 2018) or islet eQTL (right) (Varshney *et al*. 2017). Error bars denote the 95% confidence intervals. B: fGWAS conditional enrichment analysis testing the contribution of islet TC or ATAC-seq peak annotations after conditioning on histone-only based annotations such as active TSS and active enhancer chromatin states in islets. C: Maximum SNP PPA per T2D (BMI-unadjusted) GWAS locus after functional reweighting using a model with islet chromatin states and ATAC-seq peak annotations (x axis) or chromatin states, ATAC-seq peaks and TC (y axis) annotations. D: The LCORL T2D GWAS locus showing SNPs in the 99% credible set from genetic fine-mapping. This locus comprises genes *DCAF16, NCAPG and LCORL*. The lead GWAS SNP is labelled in red, along with LD r^2^ > 0.8 proxy SNPs in the top track. Also shown are CAGE, TC, ATAC-seq and chromatin state tracks. E: Browser shot of the *DCAF16* and *NCAPG* promoter regions where rs7667864 and eQTL lead SNP rs2074974 overlap an ATAC-seq peak. An overlapping CAGE element showed significant activity in the MPRA assay. Also shown are TF motifs with positive or negative coefficients from the MPRA lasso regression analysis.

We next asked to what extent TCs or ATAC-seq annotations add information above and beyond chromatin state annotations in the GWAS and eQTL enrichment models. We performed conditional analyses in fGWAS where, first, the enrichment parameters for active TSS or active enhancer chromatin states were modeled and fixed to their maximum likelihood values. Second, an additional parameter for either TCs or ATAC-seq peaks was estimated. We then asked if the added enrichment parameter was significant (confidence intervals above zero). We observed that TCs had a higher conditional enrichment over enhancer states for T2D (Figure 4B) compared to ATAC-seq peaks. TCs also showed a higher conditional enrichment over enhancer and promoter states for islet eQTL loci compared to ATAC-seq peaks (Figure 4B).

We then sought to utilize this new set of islet TC functional annotations to fine-map GWAS loci and reweight the SNP posterior probabilities of association (PPAs). We performed functional reweighting of T2D GWAS (BMI-unadjusted) using annotations such as chromatin states, ATAC-seq peaks and TCs with fGWAS. We observed that including functional annotations resulted in higher maximal SNP PPA at a locus when compared to maximal SNP PPAs from genetic fine mapping alone in many cases (Supplementary Figure 7), consistent with other studies (Pickrell 2014; Mahajan *et al*. 2018). We then compared functional reweighting when using chromatin states and ATAC-seq peaks without or with TCs and observed that some loci achieve a higher maximal reweighted SNP PPA when TC annotations are included in the model, suggesting that TCs add valuable information in fine-mapping (Figure 4C). We highlight one such GWAS locus named LCORL (lead SNP rs12640250, p value = 3.7×10^-8^). The 99% genetic credible set at this locus includes 74 variants (Mahajan *et al*. 2018), where the lead SNP rs12640250 obtains the maximum PPA = 0.15 (Supplementary Figure 8A). This GWAS signal co-localizes with an eQTL signal in islets for the gene *DCAF16* (lead eQTL SNP rs2074974, eQTL beta distribution-adjusted p values = 7.7×10^-4^, COLOC (Giambartolomei *et al*. 2014) q value = 5.44×10^-4^) (Viñuela *et al*. 2020). Functional reweighting using Islet chromatin states, ATAC-seq peaks and TCs resulted in 44 SNPs in the 99% credible set, where rs7667864 (genetic PPA = 0.12, LD r^2^ 0.97 with the lead GWAS SNP) obtained the maximum reweighted PPA = 0.62 (Supplementary Figure 8B). This SNP overlaps an ATAC-seq peak and a TC in islets (Figure 4D, E). The eQTL lead SNP rs2074974 (genetic PPA = 0.026, LD r2 = 0.96 with lead GWAS SNP) occurs upstream of the TC and overlaps the ATAC-seq peak, and obtains a reweighted PPA = 0.096 (Figure 4E). An element overlapping this TC showed significant activity in our MPRA assay (Z score = 18.48, p value = 1.56×10^-76^) and several TF motifs that showed positive MPRA lasso regression coefficients also occur in this region (Figure 4D). These analyses demonstrate that transcription initiation sites demarcate active regulatory elements in islets and this information can be useful in fine mapping and prioritizing variants.

## Discussion

Profiling TSSs using CAGE, we identified tens of thousands of TCs in 57 human pancreatic islet samples and present a consensus set of 9,954 reproducible TCs. Islet TCs were enriched to occur in islet promoter chromatin states and ATAC-seq peaks, which reflects the expected chromatin landscape at regions where transcription initiation occurs. Comparison of islet consensus TCs with those identified across 118 diverse tissues revealed that 20% of islet TCs were islet-specific. The islet-specific TCs occurred more distal to known protein-coding gene TSSs. Our analyses also highlighted the differences in the chromatin architecture underlying islet TC in islets compared to the whole pancreas tissue. These analyses demonstrate that islet TCs mark active, specific and relevant islet regulatory elements.

Surveying the TF footprint motifs occurring in TCs revealed that several ETS family footprint motifs were highly enriched in transcribed and accessible promoter regions, and that these motifs were also strong predictors of the elements’ activity in the MPRA assay. The regulatory potential of ETS family motifs has been described in the literature. One study demonstrated that orienting for islet eQTL SNPs occurring in ETS footprint motifs, the base preferred in the motifs was significantly more often associated with increased expression of the target gene (Viñuela *et al*. 2020). Another study utilizing an MPRA assay with tiled sequences in HepG2 and K562 cell lines also observed high regulatory activities of ETS motifs (Ernst *et al*. 2016). Our concordant findings with these studies using orthogonal datasets highlights the robustness and utility of our islet TC map. Our work also revealed that transcribed and accessible enhancer regions were most enriched to overlap TF footprint motifs for the RFX family of TFs. We previously showed that RFX footprint motifs are confluently disrupted by T2D GWAS risk alleles (Varshney *et al*. 2017), which are enriched to occur in islet-specific enhancer regions. These observations together highlight the role of islet specific enhancer regions, and the potential of ATAC-seq and CAGE profiling to pinpoint the active regulatory nucleotides within enhancer regions.

Utilizing an MPRA approach, we observed 2,279 CAGE elements showed significant enhancer activity, which represents ~68% of the testable elements and again highlights how CAGE profiling identifies active regulatory elements. It is interesting to note that our MPRA platform where the tested element is cloned downstream of the GFP gene is an ‘enhancer’ reporter assay. While MPRA vectors are episomal and do not recapitulate the underlying chromatin context, our results nevertheless show that sequences associated with native promoter chromatin state landscapes can show strong enhancer activity when cloned downstream of a reporter gene. We note that only a small fraction of CAGE TCs identified in our work (0.4%) overlapped with the active enhancer chromatin state. Studies have shown that gene distal transcripts are more unstable, which would therefore be difficult to profile from a total RNA sample. Of course, given the relative instability of enhancer RNAs, enhancer chromatin sites may be actively transcribed but fall below the limits of detection of CAGE. Therefore, it is plausible that islet CAGE profiling from total RNA, as we have performed, would comprise more stable promoter-associated RNA transcripts and have a lesser representation of weaker transcripts originating from enhancer regions. Recent new technologies such as native elongating transcript CAGE (NET-CAGE) show promise in more efficiently identifying more unstable transcripts from fixed tissues (Hirabayashi *et al*.2019). We note that CAGE-based enhancer calls represent only the most transcriptionally active subset of enhancers in the genome. The Roadmap Epigenomics Consortium used DNase-seq and histone modification ChIP-seq to identify 2,328,936 enhancers across 127 cell types (The Roadmap Epigenomics Consortium *et al*.2015) whereas, the FANTOM5 Consortium, in their extensive catalog of CAGE enhancers identified 43,011 enhancers across 808 CAGE libraries (Andersson *et al*. 2014).

In our previous work (Varshney *et al*. 2019; Viñuela *et al*. 2020), we showed that genetic variants in more cell type-specific enhancer regions have lower effects on gene expression than the variants occurring in more ubiquitous promoter regions. This finding is consistent with our observation that enhancer chromatin state regions comprise a smaller proportion of active transcription initiation sites and lower transcriptional activities relative to promoter chromatin state regions. The transcription initiation landscape could also change from the basal cell state under stimulatory conditions where relevant enhancers help orchestrate a cellular response.

Integrating our islet TC information with GWAS and eQTL data revealed the potential to better understand the mechanisms underlying these associations. We reasoned that if CAGE TCs represent active sites within regulatory elements and are functional for gene regulation and disease, T2D GWAS and islet eQTL variants would be enriched at these sites and SNPs occurring at these sites would more likely be causal. Indeed, regions supported by TCs, ATAC-seq peaks and enhancer chromatin states (transcribed, accessible enhancer regions) were most enriched to overlap T2D GWAS loci. This enrichment was higher than in regions only informed by ATAC-seq peaks and enhancer chromatin states, indicating that the small set of TCs in enhancer regions delineate highly relevant elements. Our work demonstrates that transcription initiation information profiled using CAGE in islets can be used in addition to other relevant epigenomic information such as histone mark informed chromatin states and chromatin accessibility in nominating causal variants and biological mechanisms.

## Data Availability

Raw and processed Islet CAGE data has been submitted to dBGaP (accession no. phs001188.v2.p1), raw and processed MPRA data has been submitted to GEO (GSE137693). A UCSC browser session with Islet CAGE tag clusters and tracks, chromatin states, mRNA-seq, ATAC-seq is available at http://genome.ucsc.edu/s/arushiv/cage_2020. Code to run analyses presented in the paper is shared on GitHub (https://github.com/ParkerLab/islet_cage) and the accompanying data files such as processed bam files, TCs, chromatin states etc. are shared on Zenodo (https://zenodo.org/record/3524578).

## Acknowledgements

We thank Sally A. Camper, Mats Ljungman, Cristen J. Willer and members of the Parker lab for their feedback. We acknowledge support from the University of Michigan Rackham Predoctoral Fellowship (to A.V.), T32 HG00040 from the National Human Genome Research Institute of the N.I.H. (to P.O.), ADA Accelerator grant 1-18-ACE-15 (to M.L.S.), NIH intramural support from project ZIA-HG000024 (to F.S.C.), ADA Pathway to Stop Diabetes Grant 1-14-INI-07 and the National Institute of Diabetes and Digestive and Kidney Diseases grant R01 DK117960 (to S.C.J.P.) S.C.J.P is the guarantor of this work and, as such, had full access to all the data in the study and takes responsibility for the integrity of the data and the accuracy of the data analysis.

## Materials and Methods

### Islet Procurement and Processing

Islet samples from organ donors were received from the Integrated Islet Distribution Program, the National Disease Research Interchange (NDRI), and Prodo-Labs. Islets were shipped overnight from the distribution centers. Upon receipt, we prewarmed islets to 37 °C in shipping media for 1–2 h before harvest; ~2,500–5,000 islet equivalents (IEQs) from each organ donor were harvested for RNA isolation. We transferred 500–1,000 IEQs to tissue culture-treated flasks and cultured them as in the work in (Gershengorn *et al*. 2004).

### RNA isolation, CAGE-seq library preparation and sequencing

Total RNA from 2000-3000 islet equivalents (IEQ) was extracted and purified using Trizol (Life Technologies). RNA quality was confirmed with Bioanalyzer 2100 (Agilent); samples with RNA integrity number (RIN) > 6.5 were prepared for CAGE sequencing. 1ug Total RNA samples were sent to DNAFORM, Japan where CAGE libraries were generated. The library preparation included polyA negative selection and size selection (<1000bp) to attempt to enrich for the short and non-polyadenylated eRNA transcripts. Stranded CAGE libraries were generated for each islet sample using the no-amplification non-tagging CAGE libraries for Illumina next-generation sequencers (nAnT-iCAGE) protocol (Murata *et al*. 2014). Each islet CAGE library was barcoded, pooled into 24-sample batches, and sequenced over multiple lanes of HiSeq 2000 to obtain paired-end 126 bp sequences. All procedures followed ethical guidelines at the National Institutes of Health (NIH).

### CAGE data mapping and processing

We processed islet CAGE data uniformly with CAGE data for other tissues included in separate ongoing projects. Because read lengths differed across libraries, we trimmed all reads to 51 bp, which was the minimum read length for all libraries, using fastx_trimmer (FASTX Toolkit v. 0.0.14). Adapters and technical sequences were trimmed using trimmomatic (v. 0.38; paired-end mode, with options ILLUMINACLIP:adapters.fa:1:30:7:1:true). To remove potential contamination, we mapped to the *E. coli* chromosome (genome assembly GCA_000005845.2) with bwa mem (v. 0.7.15; options: -M). We then removed read pairs that mapped in a proper pair (with mapq >= 10) to *E. coli*. We mapped the remaining reads to hg19 using STAR (v. 2.5.4b; default parameters) (Dobin *et al*. 2013). We pruned the mapped reads to high quality autosomal read pairs (using samtools view v. 1.3.1; options -f 3 -F 4 -F 8 -F 256 -F 2048 -q 255)(Li *et al*. 2009). We then performed UMI-based deduplication using umitools dedup (v. 0.5.5; --method directional).

We selected 57 islet samples with strandedness measures >0.85 calculated from QoRTS (Hartley and Mullikin 2015) for all downstream analyses. As another quality measure for our data, we compared overlap enrichment of CAGE TCs in chromatin states for islets and other FANTOM tissues for which chromatin state data was also publicly available. We saw that in our islet data and in FANTOM tissues, CAGE TCs are overwhelmingly enriched to overlap active TSS chromatin states, indicating that data from our protocol are comparable to existing data (Supplementary figure 9).

### Tag cluster identification

We used the paralu method to identify clusters of CAGE start sites (CAGE tag clusters) (Frith *et al*. 2008). The algorithm uses a density parameter d and identifies segments that maximize the value of (Number of events - d * size of the segment (bp)). Here, large values of d would favor small, dense clusters and small values of d would favor larger more rarefied clusters. The method identifies segments over all values of d beginning at the largest scale, where d = 0, where all of the events are merged into one big cluster. It then calculates the density (events per nucleotide) of every prefix and suffix of the big cluster. The lowest value among all of these densities is the maximum value of d for the big cluster because at higher values of d the big cluster will no longer be a maximal-scoring segment (because zero-scoring prefixes or suffixes are not allowed).

We called TCs in each individual sample using raw tag counts, requiring at least 2 tags at each included start site and allowing single base-pair tag clusters (‘singletons’) if supported by >2 tags. We then merged the tag clusters on each strand across samples. For each resulting segment, we calculated the number of islet samples in which TCs overlapped the segment. We included the segment in the consensus TCs set if it was supported by independent TCs in at least 10 individual islet samples. This threshold was selected based on comparing the number of tag clusters with the number of samples across which support was required to consider the segment (Supplementary Figure 1). We then filtered out regions blacklisted by the ENCODE consortium due to poor mappability (wgEncodeDacMapabilityConsensusExcludable.bed and wgEncodeDukeMapabilityRegionsExcludable.bed) using bedtools subtract to obtain the final set of Islet tag cluster regions used in all downstream analyses.

### FANTOM CAGE datasets

We obtained the set of ‘robust CAGE peaks’ identified by the FANTOM 5 consortium (The FANTOM Consortium *et al*. 2014) using CAGE libraries (CAGE sequencing on HeliScope Single Molecule Sequencer (hCAGE)) of 988 human cell lines or tissues (http://fantom.gsc.riken.jp/5/datafiles/latest/extra/CAGE_peaks/hg19.cage_peak_phase1and2combined_coord.bed.gz). These peaks were identified using the decomposition-based peak identification (DPI) method (The FANTOM Consortium *et al*. 2014), followed by filtering ‘robust’ peaks that included a CAGE tag (TSS) with more than 10 read counts in at least 1 sample and 1 tags per million (TPM). For a more direct comparison of islet TCs with TCs from other tissues, we downloaded the CAGE transcription start site (CTSS) data for 118 tissue types (from http://fantom.gsc.riken.jp/5/datafiles/latest/basic/human.tissue.hCAGE/) and called tag clusters for each tissue sample using the paraclu method (Frith *et al*. 2008) as described above, with the same parameters.

### Chromatin state analysis

We generated ChromHMM states in islets along with 30 other tissues previously (Varshney et al. 2017) However, since the statistical power to identify enriched regions depends on the ChIP-seq read depth, in this previous study we uniformly downsampled the ChIP seq datasets to the depth of 20 million reads across the 31 tissues. This allowed comparable chromatin state territories across tissues and ensured that chromatin state territories were not heavily driven by high sequencing depth. Other studies have presented independent chromHMM states in islets, however, there isn’t a consensus set of states (Thurner et al. 2018; Miguel-Escalada et al. 2019). For the current manuscript, we aimed to compare islets and three other relevant tissues (Skeletal Muscle, Liver and Adipose). To maximize power to detect enriched states in this smaller subset of tissues in our study, we downsampled read depth to the mean depth across the four tissues within each mark and then generated harmonized ChromHMM states across the tissue types. These subsampling depths were H3K27ac = 28,123,470, H3K27me3 = 53,603,910, H3K36me3 = 63,763,910, H3K4me1 = 63,441,280 and H3K4me3 = 62,220,940. We list references for the ChIP-seq datasets we utilized in Supplementary Table 5. We performed read mapping and integrative chromatin-state analyses in a manner similar to that of our previous reports (Scott *et al*. 2016; Varshney *et al*. 2017) and followed quality control procedures reported by the Roadmap Epigenomics Study (The Roadmap Epigenomics Consortium *et al*.2015). Briefly, we trimmed reads across datasets to 36bp and overrepresented adapter sequences as shown by FASTQC (version v0.11.5) using cutadapt (version 1.12) (Martin 2011). We then mapped reads using BWA (version 0.5.8c), removed duplicates using samtools (Li *et al*. 2009), and filtered for mapping quality score of at least 30. To assess the quality of each dataset, we performed strand cross-correlation analysis using phantompeakqualtools (v2.0; code.google.com/p/phantompeakqualtools) (Landt *et al*. 2012). We converted bam files for each dataset to bed using the bamToBed tool, followed by randomly subsampling each dataset bed file to the thresholds mentioned above. Chromatin states were learned jointly for the three cell types using the ChromHMM (version 1.10) hidden Markov model algorithm at 200-bp resolution to five chromatin marks and input (Ernst and Kellis 2010, 2012; Ernst *et al*. 2011). We ran ChromHMM with a range of possible states and selected a 11-state model, because it most accurately captured information from higher-state models and provided sufficient resolution to identify biologically meaningful patterns in a reproducible way. To assign names to chromatin states that are consistent with previously published states, we performed enrichment analyses in ChromHMM comparing our states with the states reported previously (Varshney *et al*. 2017) for the four matched tissues. We assigned each state with the state name that was most strongly enriched to overlap that state.

### ATAC-seq data analysis

We used previously published chromatin accessibility data profiled using ATAC-seq in islets from two human organ donor samples (Varshney *et al*. 2017). For each sample, we trimmed reads to 36 bp (to uniformly process ATAC-seq from other tissues for ongoing projects) and removed adapter sequences using Cutadapt (version 1.12) (Martin 2011), mapped to hg19 used bwa-mem (version 0.7.15-r1140) (Li 2013), removed duplicates using Picard (http://broadinstitute.github.io/picard) and filtered out regions blacklisted by the ENCODE consortium due to poor mappability (wgEncodeDacMapabilityConsensusExcludable.bed and wgEncodeDukeMapabilityRegionsExcludable.bed). For each tissue we subsampled both samples to the same depth (18M reads for Islet samples) so that each tissue had overall similar genomic region called as peaks. We used MACS2 (https://github.com/taoliu/MACS), version 2.1.0, with flags “-g hs–nomodel–shift -100–extsize 200 -B–broad–keep-dup all,” to call peaks and retained all broad-peaks that satisfied a 1% FDR.

### Overlap enrichment between TCs and annotations

We calculated the enrichment for Islet TCs to overlap annotations such as different Islet chromatin states, Islet ATAC-seq peaks and various ‘common’ annotations. Common annotations imply annotations that don’t vary across cell types such as coding gene regions, intronic regions or annotations created by merging epigenomic data such as histone modification peaks across cell types. We utilized 29 total static annotation bed files supplied by (Finucane *et al*. 2015) (https://data.broadinstitute.org/alkesgroup/LDSCORE/baseline_bedfiles.tgz). These included coding, untranslated regions (UTRs), promoter and intronic regions obtained from UCSC (Kent *et al*. 2002); the histone marks monomethylation (H3K4me1) and trimethylation (H3K4me3) of histone H3 at lysine 4 and acetylation of histone H3 at lysine 9 (H3K9ac) (The ENCODE project Consortium 2012; Trynka *et al*. 2013; The Roadmap Epigenomics Consortium *et al*. 2015) and acetylation of histone H3 at lysine 27 (H3K27ac) (Hnisz *et al*. 2013; Schizophrenia Working Group of the Psychiatric Genomics Consortium 2014); open chromatin, as reflected by DNase I hypersensitivity sites (DHSs) (Trynka *et al*. 2013; Gusev *et al*. 2014); combined chromHMM and Segway predictions(Hoffman *et al*. 2013), which partition the genome based on distinct and recurring patterns of histone marks into seven underlying chromatin states; regions that are conserved in mammals(Lindblad-Toh *et al*. 2011; Ward and Kellis 2012); super-enhancers, which are large clusters of highly active enhancers(Hnisz *et al*. 2013); and enhancers with balanced bidirectional capped transcripts identified using CAGE in the FANTOM5 panel of samples, (called Enhancer (Andersson)) (Andersson *et al*. 2014). Histone marks included in the static annotation set included merged histone mark data from different cell types into a single annotation.

Enrichment for overlap between each Islet tag clusters and regulatory annotations was calculated using the Genomic Association Tester (GAT) tool (Heger *et al*. 2013). To ask if two sets of regulatory annotations overlap more than that expected by chance, GAT randomly samples segments of one regulatory annotation set from the genomic workspace (hg19 chromosomes) and computes the expected overlaps with the second regulatory annotation set. We used 10,000 GAT samplings for each enrichment run. GAT outputs the observed overlap between segments and annotation along with the expected overlap and an empirical p-value.

### Aggregate signal

We generated the ATAC-seq density plot over islet TC midpoints using the Agplus tool (version 1.0) (Maehara and Ohkawa 2015). We used the ATAC-seq signal track for reads per 10 Million to aggregate over stranded TCs.

To obtain CAGE tracks, we merged CAGE bam files for islet samples that passed QC (see CAGE data processing section) and obtained the read 1 start sites or TSSs. To better visualise the CAGE signal, we then flanked each TSS 10bp upstream and downstream and normalized the TSS counts to 10M mapped reads. We generated CAGE density plots over ATAC-seq narrow peak summits by using the agplus tool.

To obtain aggregate CAGE signal over TF footprint motifs, we oriented the CAGE signal with respect to the footprint taken on the plus strand. We used HTSeq GenomicPosition method (Anders *et al*. 2015) to obtain the sum of CAGE signal at each base pair relative to the footprint motif mid point.

### Enrichment for Islet TF footprint motifs to overlap TC-related annotations

We compared the enrichment of Islet TF footprint motifs in several TC-related annotations such as upstream and downstream 500bp regions of TCs and Islet TCs that occurred in accessible enhancer states vs those that occurred in accessible promoter states using the GAT tool similarly as described above (Heger *et al*. 2013). TF footprint motifs are occurrences of TFs motifs (obtained from databases of DNA binding motifs for several TFs) in accessible chromatin regions (identified from assays such as ATAC-seq). We utilized previously published islet TF footprint motifs (Varshney *et al*. 2017), which were generated using ATAC-seq data in two islet samples and DNA binding motif information for 1,995 publicly available TF motifs (Jolma *et al*. 2013; Kheradpour and Kellis 2014; Mathelier *et al*. 2016).

We obtained the 500bp upstream or downstream regions of each TC (upstream/downstream regions were determined based on TC strand; TC region itself was not included). We generated the list of regions where islet TCs overlapped ATAC-seq peaks and any enhancer states (Active enhancer, Weak enhancer, or Genic enhancer) using BEDTools intersect - we referred to these as ‘TCs in accessible enhancers’. Similarly, we also generated the list of regions where islet TCs overlapped ATAC-seq peaks and any TSS/promoter states (Active TSS, Weak TSS, Flanking TSS) - we referred to these as ‘TCs in accessible promoters’. as segments.

In the GAT analyses, the TC-related annotations were considered as ‘annotations’ (argument -a) and footprint motif occurrences for each known motif were considered as ‘segments’ (argument -s). Since the TF footprint motifs only occur in ATAC-seq peaks, we considered these peaks as the ‘workspace’ (argument -w) to sample segments. We used 10,000 GAT samplings for each enrichment run. We accounted for the 1,995 footprint motifs being tested against each TC-related annotation by performing an FDR correction with the Benjajmini-Yekutieli method (Benjamini and Yekutieli 2001) using the stats.multitest.multipletests function from the statsmodels library in Python (Seabold and Perktold 2010). Significant enrichment was considered at 5% FDR threshold.

### Comparison of features with Roadmap chromatin states

We downloaded the chromatin state annotations identified in 127 human cell types and tissues by the Roadmap epigenomics project (The Roadmap Epigenomics Consortium *et al*. 2015) after integrating ChIP-seq data for five histone 3 lysine modifications (H3K4me3, H3K4me1, H3K36me3, H3K9me3 and H3K27me3) that are associated with promoter, enhancer, transcribed and repressed activities, across each cell type. For each TC feature, for example, TCs in ATAC-seq peaks within islet enhancer chromatin states, we identified segments occurring proximal to (within 5kb) and distal from (further than 5kb) known protein coding gene TSS (gencode V19). For each such segment, we identified the maximally overlapping chromatin state across 98 cell types publicly available from the Roadmap Epigenomics project in their 18 state ‘extended’ model using BEDtools intersect. We then ordered the segments using clustering (hclust function in R) based on the gower distance metric (daisy function in R) for the roadmap state assignments across 127 cell types.

### Experimental validation using MPRA

#### 1. Selection of CAGE elements

We generated a library of islet CAGE elements to test in the MPRA assay by using two approaches. First, we identified clusters of CAGE tags in each islet sample by simply concatenating tags that occurred within 20bp. We retained clusters with at least two tags in each islet sample. We then merged these cluster coordinates across samples and retained clusters supported by at least 15 samples, representing a highly reproducible set. Second, we complemented this approach by also including the set of FANTOM 5 ‘robust’ CAGE peaks that were also supported by CAGE tags in at least 15 samples. 94% of the selected CAGE robust peak regions were already included in the selected CAGE 20 bp clusters; we reasoned that the remaining 6% of CAGE peaks represented relevant and reproducible CAGE elements missed by the 20 bp concatenation approach. We therefore took the union of these two sets of CAGE elements and created 198 bp oligo sequences centered on each element for cloning into the MPRA vector. When a CAGE element was longer than 198 bp, we tiled 198 bp oligos over the element, offset by 100 bp. Through this approach, we included a total of 7,188 CAGE elements (each 198 bp long). We note that these CAGE elements represent slightly different coordinates from the TCs coordinates presented elsewhere in the paper that were identified using the paraclu method. While the paraclu approach of calling TCs was adopted after the MPRA experiments were already performed, 6,810 (94.7%) of the CAGE elements included in MPRA experiment overlapped the final set of TCs presented in the manuscript.

We synthesized 230-bp oligos (198bp CAGE element flanked by 16bp anchor sequences) (Agilent Technologies). We PCR-amplified oligos to add homology arms for Gibson assembly cloning, and gel-purified the PCR products (i.e. inserts) using the Zymoclean Gel DNA Recovery Kit (Zymo Research). We used the NEBuilder HiFi DNA Assembly Kit (NEB) to assemble the purified inserts and the backbone of MPRA plasmid (previously digested with EcoRV). After column purification of the reaction using the DNA Clean and Concentrator-5 kit (Zymo), we transformed it into 10beta electrocompetent cells (NEB), and obtained 1.39 million unique transformants.

We post-barcoded the library by first digesting the library with PmeI, and then by setting up Gibson assembly to insert 16-bp random nucleotides (‘barcodes’) at the PmeI restriction site. After column purification, we transformed the reaction into electrocompetent cells (NEB), and obtained 1.44 million unique transformants. We prepared the library plasmid for transfection using the ZymoPURE Plasmid Maxiprep Kit (Zymo).

#### 2. Electroporation, RNA isolation and cDNA synthesis

We electroporated 50 ug of barcoded MPRA library into 25 million the 832/13 rat insulinoma cell line for each biological replicate (3 replicates), and harvested the cells twenty-four hours later. We isolated total RNA using TRIZOL reagent (Life Technologies) following the manufacturer’s protocol up to phase separation. After phase separation, we transferred the aqueous phase of the solution to a new 1.5 mL Eppendorf tube, added 1:1 volume of 100% ethanol, and then column-purified using the Direct-zol RNA Miniprep Kit (Zymo Research Corporation, Irvine, CA) following the manufacturer’s protocol. We further purified mRNA using Dynabeads oligo(dT) beads (Thermo Fisher, Carlsbad, CA). We treated 2 ug of mRNA with RNase-free DNaseI (Invitrogen, Carlsbad CA) to eliminate possible plasmid DNA contamination, and then reverse transcribed 1ug into cDNA using the SuperScript III First-Strand Synthesis kit (Invitrogen) with a custom primer that specifically recognizes ‘MPRA transcripts’ (i.e. mRNA that had been transcribed from the MPRA plasmids). The other 1ug of mRNA was used in a enzyme-negative reaction to determine DNA/plasmid contamination in cDNA, which we did not. To eliminate any residual plasmid contamination in cDNA samples, we treated cDNA with DpnI (NEB), and purified the reaction using the DNA Clean and Concentrator-5 kit (Zymo).

#### 3. Construction of Illumina sequencing libraries

We constructed Illumina sequencing library via two serial rounds of PCR. In the first round, we used a primer set to specifically amplify STARR transcripts using cDNA as starting material. In the second round, we used a primer set to append the P5/P7 Illumina sequences using the PCR product from the first round as starting material. In both rounds, we PCR-amplified the fragments until the amplification curve reached a mid-log phase, and then purified the products for subsequent steps using the DNA Clean and Concentrator-5 kit (Zymo).

#### 4. CAGE element-barcode pairing

To identify the barcodes corresponding to each CAGE element in the MPRA plasmid, 1ng of each library constructed was used in a polymerase chain reaction with primers flanking the allele and the barcode to generate fragments which were subsequently gel verified and extracted using Zymo gel extraction kit (Zymo). 25ng of the purified product was subjected to self-ligation at 16 °C overnight in a total volume of 50uls and column purified using Qiagen (Qiagen) PCR purification kit as per manufacturers recommendations. The purified fragments were subsequently treated with 10U of Plasmid-Safe ATP-Dependent DNase for 1 hour in the presence of 25mM ATP to remove unligated linear DNA fragments. 1ul of the recircularized fragments were subjected to another round of PCR resulting in a smaller fragment. Briefly an aliquot of this product was diluted 1:10 in DNAse/RNase free water and amplified in a PCR reaction, with Illumina P5 and P7 adapters, until saturation to generate libraries. The libraries were subsequently column purified, quantified and sequenced.

#### 5. Data analysis

The MPRA barcode sequencing data included the input DNA barcode library along with three cDNA barcode libraries representing three biological replicated. We processed this data through a custom pipeline which quantified barcode counts while accounting for sequencing errors. We extracted the DNA barcodes from the input DNA library (first 16 bp of the read-1 fastq file) and clustered these at an edit distance of 0 followed by computing the DNA counts for each read group of DNA sequencing files. We then aggregated the read groups and collapsed counts for barcodes using the sequence clustering algorithm Starcode (https://github.com/gui11aume/starcode) (Zorita *et al*. 2015). We repeated this process for the cDNA barcode counts for each replicate, with the added step of removing PCR-duplicated barcodes using the UMI information (UMI sequence was the reverse complement of the first 6bp of the read-2 fastq file). The pipeline is shared at https://github.com/ParkerLab/STARR-seq-Analysis-Pipeline.

We matched the barcodes with CAGE inserts using results from CAGE insert-barcode pairing experiment (data file “cage_insert_barcode_pairing.tsv” in GEO GSE137693). We first retained barcodes with at least 10 DNA counts, and further retained CAGE elements that had at least two such qualifying barcodes. This was the set of N=3,446 CAGE elements quantifiable in our assay. To quantify MPRA activities from these count-based data, we used the tool MPRAnalyze (version 1.3.1) (https://github.com/YosefLab/MPRAnalyze) (Ashuach *et al*.2019) that models DNA and RNA counts in a negative binomial generalized linear model. This approach is more robust than using metrics such as the aggregated ratio, which is the ratio of the sum of RNA counts across barcodes divided by the sum of DNA counts across barcodes and loses the statistical power provided by multiple barcodes per tested element; and the mean ratio, which is the mean of the observed RNA/DNA ratios across barcodes which can be quite sensitive to low counts and noise. We corrected for library depth for the three replicates using upper quartile normalization via the ‘estimateDepthFactors’ function in MPRAnalyze. MPRA activity is quantified by estimating the transcription rate for each element in the dataset, followed by identifying active elements that induce a higher transcription by testing against a null. MPRAnalyze fits two nested generalized linear models - the DNA model estimates plasmid copy numbers, and the RNA model is estimates transcription rate. We included barcode information in the DNA model which allows different estimated counts for each barcode, and increases the statistical power of the model. Replicate information was included in the RNA model. MPRAnalyze then tests the transcriptional activity of each element against a null distribution and computing Z and Median-Absolute-Deviation (MAD) scores. The null is based on the assumption that the mode of the distribution of transcription rate estimates is the center of the null distribution, and that values lower than the mode all belong to the null. Thus, values lower than the mode are used to estimate the variance of the null.

### Lasso regression

We used lasso regression to model TC element MPRA activity z scores as a function of TF motif occurrences within the TC elements. Lasso regression is useful when a large number of features such as hundreds of TF motifs in this case are included because it imposes a constraint on the model parameters causing regression coefficients for some variables to shrink toward zero. Features with non-zero regression coefficients are most strongly associated with the response variable.

We utilized a set of 1,995 TF motifs including their position weight matrices (PWMs), available from ENCODE, JASPAR and Jolma datasets (Jolma *et al*. 2013; Kheradpour and Kellis 2014; Mathelier *et al*. 2016), which we have also used previously (Scott *et al*. 2016; Varshney *et al*. 2017). In order to reduce motif redundancy, we performed PWM clustering in our motif database using the matrix-clustering tool from RSAT (Castro-Mondragon *et al*. 2017), with parameters -lth cor 0.7 -lth Ncor 0.7. For each of the 540 clusters obtained, we retained the motif with the highest total PWM information content. Because MPRA is an episomal assay and doesn’t recapitulate the native chromatin context, we quantified overlaps of each TC element with sequence motif scans rather than ATAC-seq informed footprint motifs. We scanned each of these motifs on the hg19 reference using FIMO (Grant *et al*. 2011). We used the nucleotide frequencies from the hg19 reference and the default p value cutoff of 10^-4^.

To quantify motif occurrences within each TC element, we considered the −log10(P value) of each motif occurrence from FIMO. Since the FIMO motif scan p-values depend on the motif length and information content etc., these log transformed P values are not directly comparable across motifs. We therefore inverse normalized the −log10(P values) for occurrences of each motif using the RNOmni package (version 0.7.1) to obtain motif ‘scores’ on a comparable normal scale. P value = 1 was included for each motif to obtain the score corresponding to no motif occurrence on the transformed scale. For each TF motif, we aligned the hg19 scan occurrences with each islet TC MPRA element using BedTools intersect and recorded the corresponding motif scores. We added the scores for TC elements that overlapped multiple occurrences for the said motif. We again inverse normalized the motif overlap score vector across the input CAGE elements for each TF motif so that the regression coefficients could be comparable across motifs. The lasso regression was run using the glmnet package (version 2.0-16) with default parameters (specifically, alpha=1, which corresponds to the lasso regression). Lambda was determined automatically by glmnet as the lambda that belonged to the model with the lowest mean cross validated error.

### 1. fGWAS analyses and fine-mapping

We used the fGWAS (version 0.3.6) (Pickrell 2014) tool to compute enrichment of GWAS and islet eQTL data in TC-related annotations along with computing conditional enrichment and fine mapping analyses. fGWAS employs a Bayesian hierarchical model to determine shared properties of loci affecting a trait. The model uses association summary level data, divides the genome into windows generally larger than the expected LD patterns in the population. The method assumes that there is either a single causal SNP in a window or none. The model defines the prior probabilities that an association lies in a genomic window and that a SNP within it is causal. These probabilities are allowed to depend on genomic annotations, and are estimated based on enrichment patterns of annotations across the genome using a Bayes approach.

We obtained publicly available summary data for T2D GWAS (Mahajan *et al*. 2018) and islet eQTL (Varshney *et al*. 2017) and organized it in the format required by fGWAS. We used fGWAS with default parameters for enrichment analyses for individual annotations in Figure 4 A. For each individual annotation, the model provided the maximum likelihood enrichment parameter. Annotations were considered as significantly enriched if the log2(parameter estimate) and respective 95% confidence intervals were above zero or significantly depleted if the log2(parameter estimate) and respective 95% confidence intervals were below zero. We performed conditional analyses using the ‘-cond’ option.

To reweight GWAS summary data based on functional annotation overlap, we used the ‘-print’ option in addition in fGWAS while including multiple annotations in the model that were individually significantly enriched or depleted. We included active TSS, active enhancer, quiescent and polycomb repressed annotations and ATAC-seq peaks with or without or TCs. To compare with T2D genetic credible sets (Mahajan *et al*. 2018), we created 1Mb windows centered on the lead variant at each primary GWAS signal. We then partitioned the rest of the genome into 1Mb windows as well. Specifying these regions using the --bed option in fGWAS enabled constraining each primary signal in a single window. The model derived enrichment priors to evaluate both the significance and functional impact of associated variants in GWAS regions, such that variants overlapping more enriched annotations carried extra weight.

