## Supplementary figures for "A transcription start site map in human pancreatic islets reveals functional regulatory signatures"

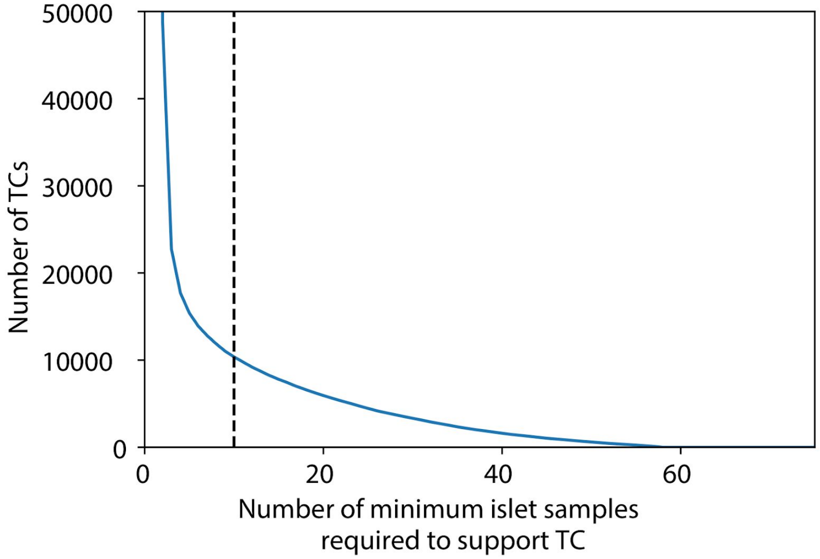


Supplementary Figure 1: Islet TC identification using CAGE data across multiple samples. TC segments called using the paraclu method in each of the 57 selected islet samples were merged in a strand specific manner. Shown here is the number of merged TC segments that overlap TCs in x or more islet samples. We required TC overlap in a minimum of 10 islet samples to include a segment in the aggregate list of islet TCs.

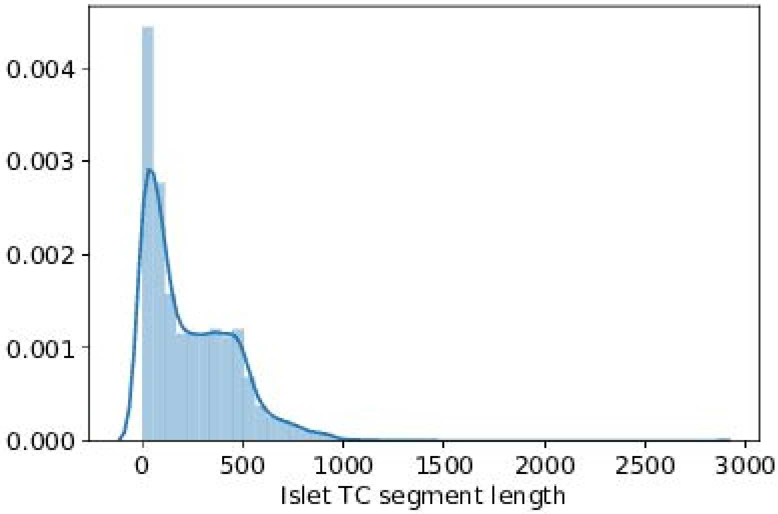


Supplementary Figure 2: Distribution of islet TC lengths.

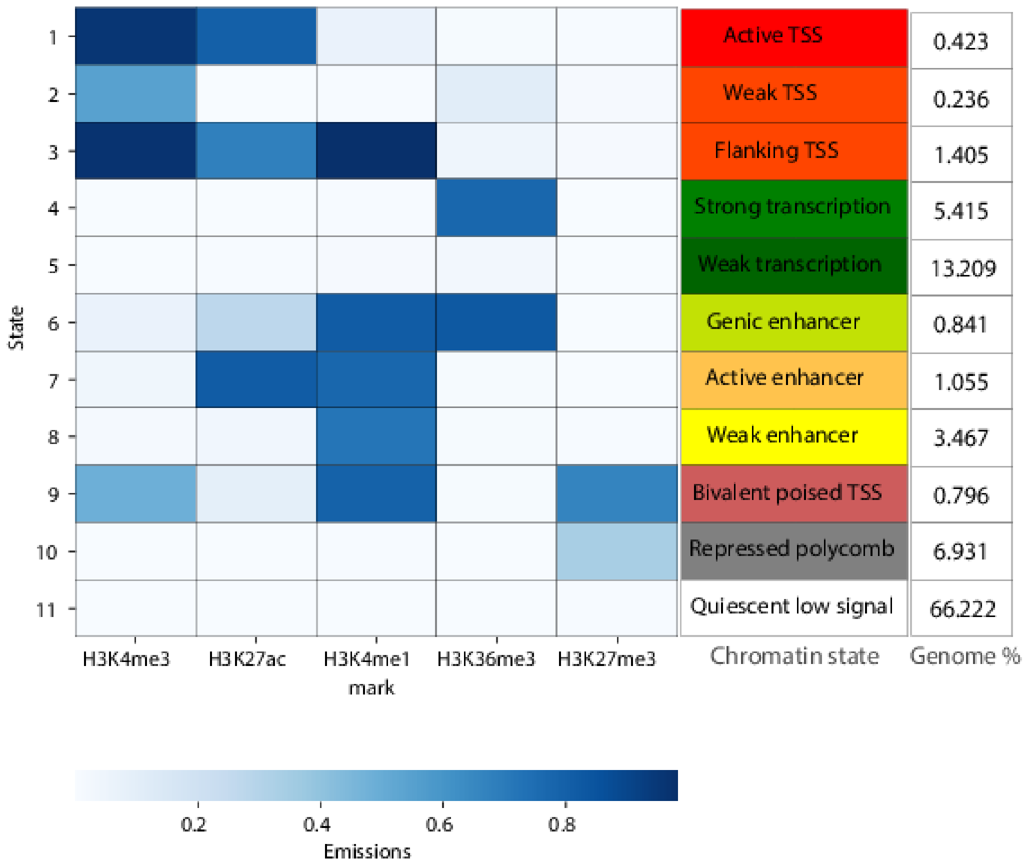


Supplementary Figure 3: 11 chromatin state model. Shown are the emission probabilities of each of the five histone marks, chromatin state annotation and the percent genome coverage of each state

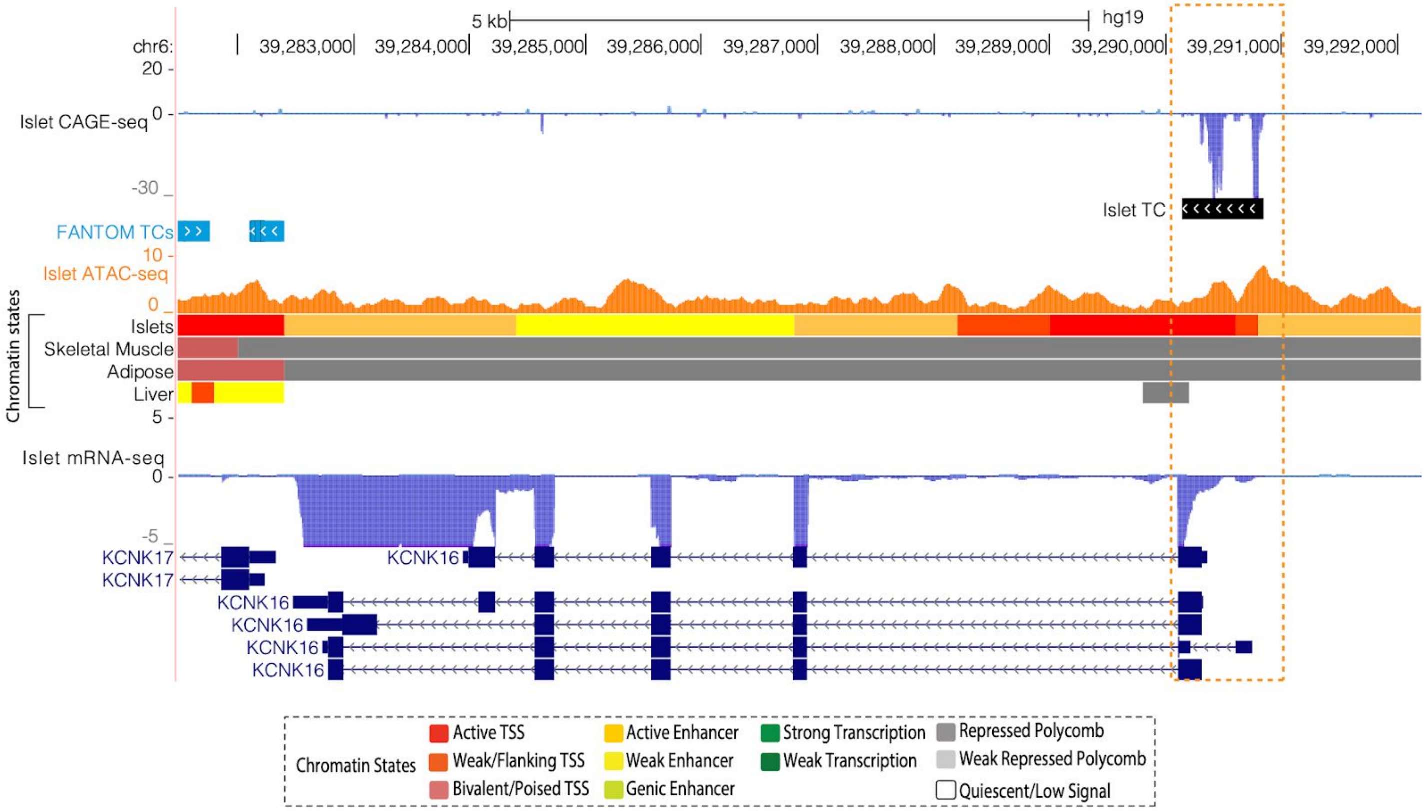


Supplementary Figure 4: *KCNK16* gene locus. Shown are tracks for Islet TCs along with TCs called across 118 FANTOM tissues.

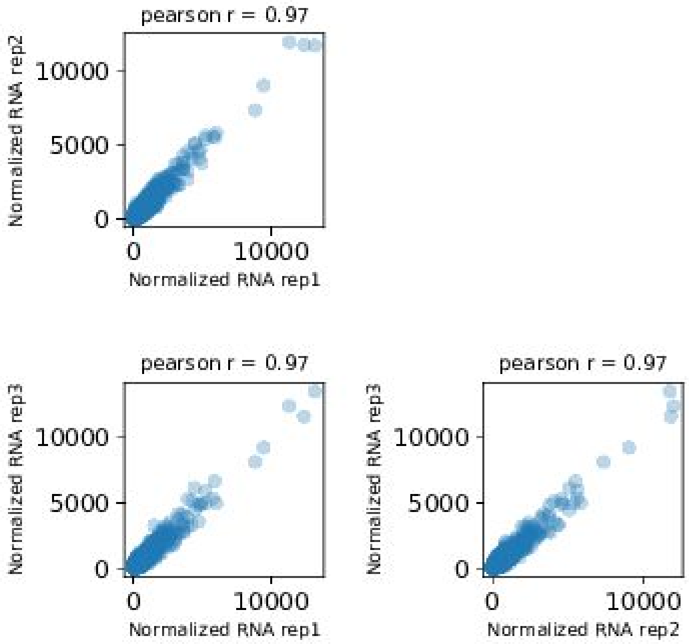


Supplementary Figure 5. Correlation between replicates for normalized total RNA counts for CAGE inserts.

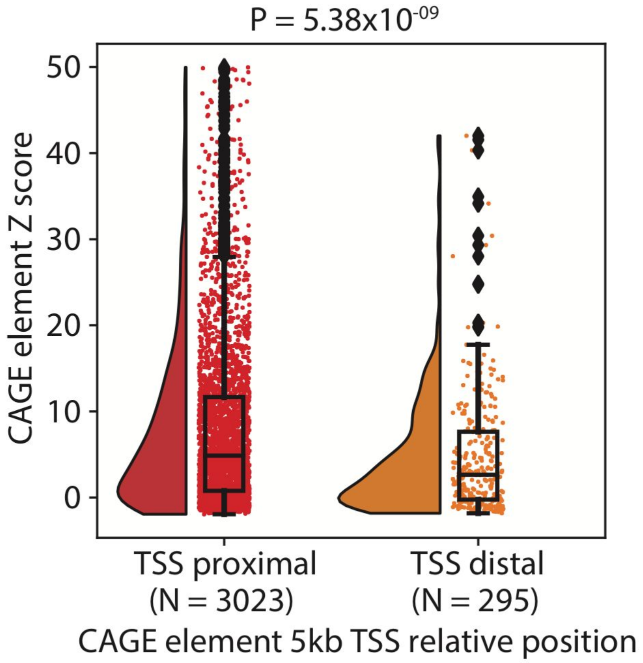


Supplementary Figure 6: MPRA activity Z scores for CAGE elements based on position relative to known protein coding gene TSSs (5kb TSS proximal or distal, Gencode V19).

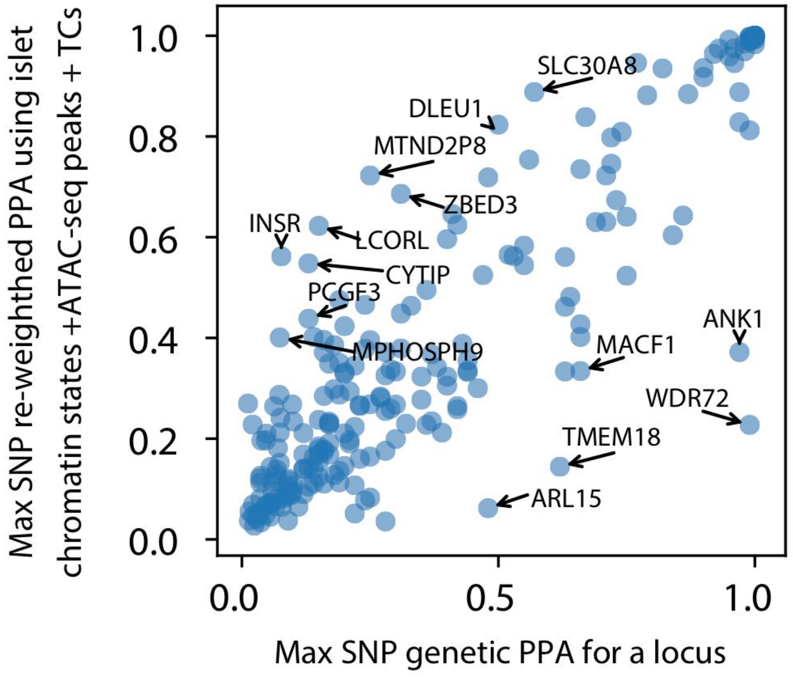


Supplementary Figure 7: The maximal SNP PPA from genetic fine-mapping for a locus vs the maximal SNP PPA after functional re-weighting using islet chromatin states, ATAC-seq peaks and TC annotations.

A

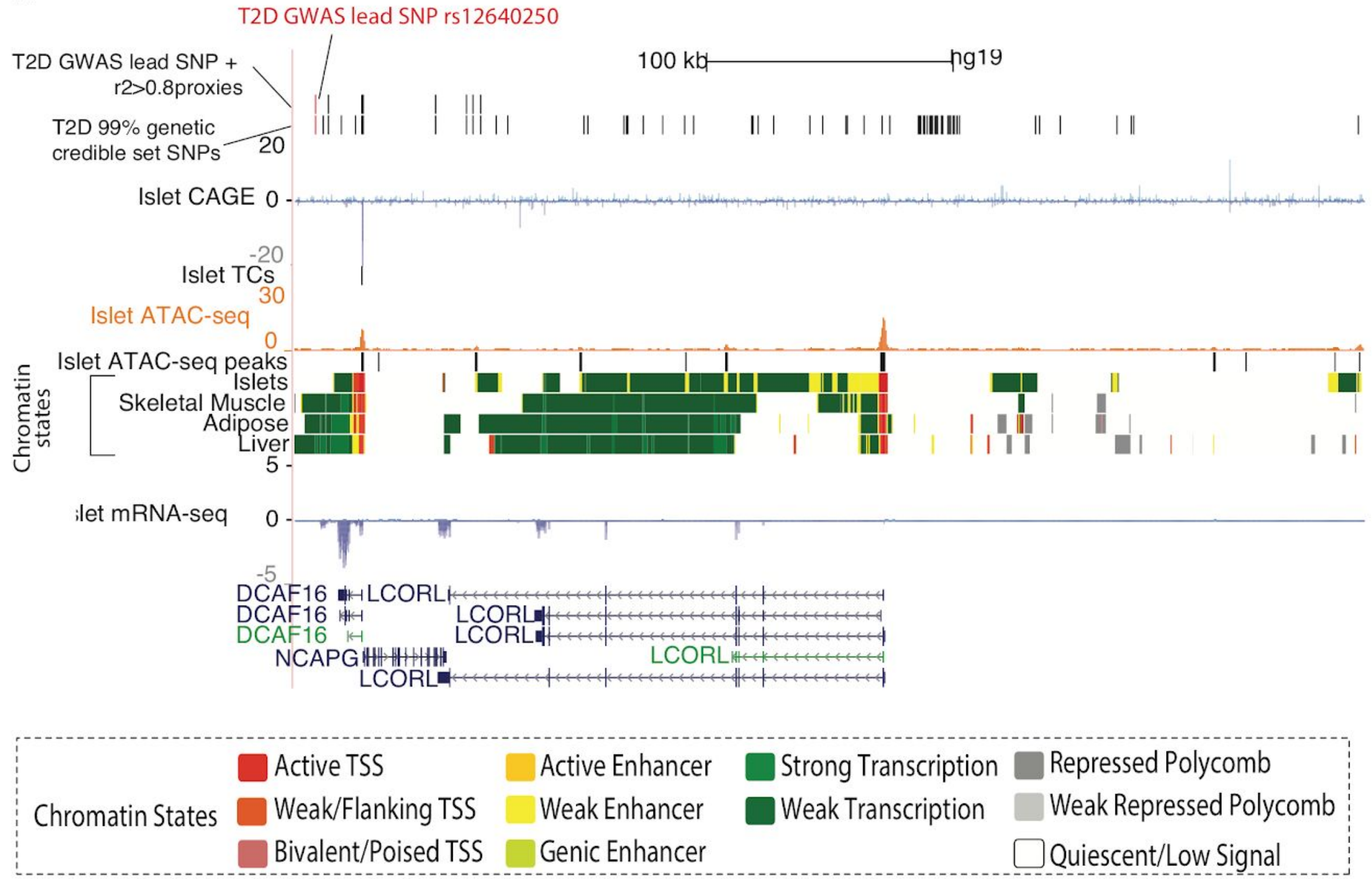

T2D GWAS lead SNP rs12640250
T2D GWAS lead SNP + r2>0.8proxies
T2D 99% genetic credible set SNPs
100 kb
hg19
20
Islet CAGE
0
-20
Islet TCs
30
Islet ATAC-seq
0
Islet ATAC-seq peaks
Chromatin states
Islets
Skeletal Muscle
Adipose
Liver
5
islet mRNA-seq
0
-5
DCAF16
DCAF16
DCAF16
NCAPG
LCORL
LCORL
LCORL
LCORL
LCORL
Chromatin States
Active TSS
Weak/Flanking TSS
Bivalent/Poised TSS
Active Enhancer
Weak Enhancer
Genic Enhancer
Strong Transcription
Weak Transcription
Repressed Polycomb
Weak Repressed Polycomb
Quiescent/Low Signal


B

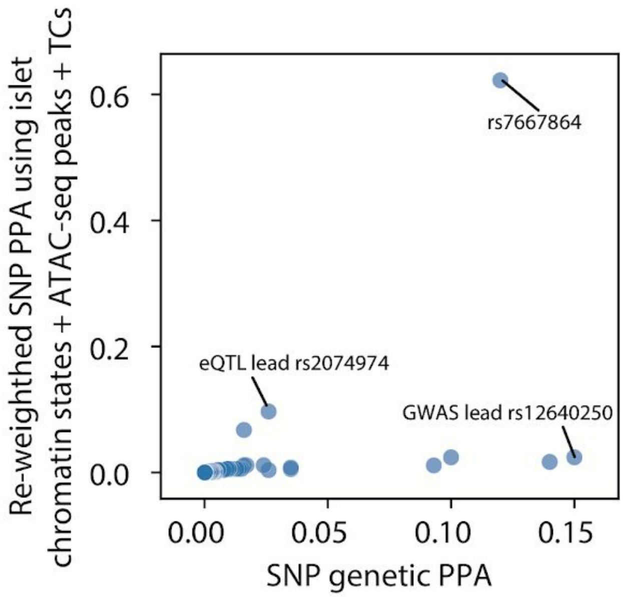

Re-weighted SNP PPA using islet chromatin states + ATAC-seq peaks + TCs
0.6
0.4
0.2
0.0
rs7667864
eQTL lead rs2074974
GWAS lead rs12640250
0.00
0.05
0.10
0.15
SNP genetic PPA

Supplementary figure 8: A: LCORL GWAS locus showing all SNPs in the 99% credible set after genetic fine-mapping. B: Genetic fine-mapping PPA and functionally re-weighted PPA for SNPs at the LCORL locus.

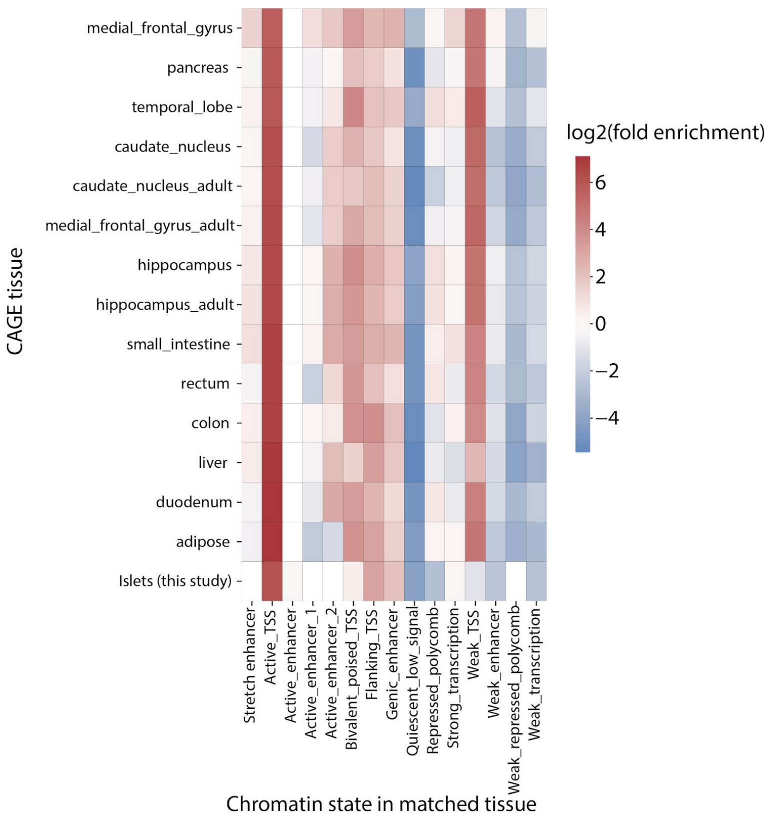


Supplementary figure 9: Overlap enrichment of CAGE TCs in various FANTOM tissue to overlap chromatin states annotations in publicly available matched tissues.
